# *Pectobacterium versatile* β-lactamase, a common good of the soft rot *Pectobacteriaceae* species complex

**DOI:** 10.1101/2025.01.30.635659

**Authors:** Camille Lorang, Pierre-Yves Canto, Erwan Gueguen, Jacques Pédron, Marie-Anne Barny

## Abstract

Little is known about the role of antibiotics and associated resistance in microbial ecosystems in the absence of clinical antibiotic pressure. The Soft Rot Pectobacteriaceae (SRP) species complex, which comprises 37 bacterial species that are collectively responsible for the severe rotting of many crops, is an interesting model to analyse the role of β-lactam and β-lactamases in natural ecosystems. In particular, within this complex, most *Pectobacterium versatile* strains harbour a β-lactamase called Bla_PEC-1_. The aim of our work was to analyse the role of Bla_PEC-1_ during infection. To this end, two *bla*_PEC-1_-deleted strains were constructed and compared with their wild-type counterparts *in vitro* and in potato tuber infections. *In vitro*, the Bla_PEC-1_ β-lactamase enables *P. versatile* to resist ampicillin or carbapenem produced by *Pectobacterium brasiliense*. In mono-infections on potato tubers, *bla*_PEC-1_-deleted strains were unaffected in terms of virulence, fitness or association with commensal bacteria. In mixed infections, the Bla_PEC-1_ β-lactamase proved necessary for the coexistence of *P. versatile* with the carbapenem-producing strain, and also for the protection of carbapenem-sensitive strains both *in vitro* and *in planta*. Interestingly, *in planta* protection was observed even if the *bla*_PEC-1_ gene was repressed and bacteria expressing Bla_PEC-1_ were in the minority within the symptom. These results indicate that Bla_PEC-1_ exerts a true β-lactamase function during the infection process and acts as a public good of the SRP species complex. Finally, our results highlight the important role of β-lactamase in maintaining of strains diversity in natural ecosystem.

**Statements relating to our ethics and integrity policies:** The data shown in this paper are available within the article and supplementary materials. The funding of the ANR, ANR-19-CE35-0016-03, is acknowledged. The funders had no role in study design, data collection and interpretation, or the decision to submit the work for publication. The authors declare that there are no conflicts of interest. All applicable local, national and international regulations and conventions, as well as normal scientific ethical practices, were followed in the preparation of this work. This manuscript has not been previously or simultaneously published or submitted elsewhere and was critically reviewed and approved by all co-authors before submission. The CRediT of all authors is provided.

## Introduction

Antibiotics are probably one of the most successful forms of chemotherapy in the history of medicine. The β-lactam antibiotics, which disrupt bacterial cell wall formation in both Gram-negative and Gram-positive bacteria, are currently the most widely used class of antibacterial agents in the treatment of infectious diseases (Bush and Bradford, 2016). All β-lactam antibiotics share the β-lactam ring as the essential core structure and are classified into four classes: penicillins, cephalosporins, carbapenems and monobactams (Kim *et al*., 2023). The massive use of β-lactams in human and animal health is leading to the constant emergence of bacterial resistance. In particular, the regular emergence of new β-lactamases that cleave and inactivate different β-lactam rings is a cause for concern (Tooke *et al*., 2019). To date, more than 2,700 unique β-lactamases have been described, they are classified into four molecular classes (classes A to D) based on their primary structure, which are further subdivided into functional groups based on their substrate preferences (Bush, 2018). For example, carbapenems are not hydrolysed by most β-lactamases, with the exception of class B and class D2df β-lactamases. It is recognized that β-lactamases are ancient enzymes that existed without the pressure of therapeutic antibiotics, and that they have emerged in clinically pathogenic species following transfer from environmental sources. The rapid spread of transmissible β-lactamases carried on plasmids or transposons that hydrolyze recently approved cephalosporins, monobactams and carbapenems is important in Gram-negative clinical pathogens (Cantón *et al*., 2008). However, the wealth of data on clinical antibiotics, resistance mechanisms and spread of resistance through mobile genetic elements contrasts with the lack of knowledge about the evolutionary and ecological processes that take place in microbial ecosystems in the absence of antibiotic pressure. In natural ecosystems, antibiotics are not present at therapeutic dose and some authors have proposed that antibiotics act as regulatory or signaling molecules rather than weapons in this context (Romero *et al*., 2011). In this scenario, β-lactamases in nature could disrupt such regulatory role or signaling. This scenario is supported by *in vitro* analysis of antibiotic role at sub-inhibitory concentration but is scarcely supported by experimental data *in natura*. To better understand the role and function of β-lactam and β-lactamases in natural ecosystems, it is important to develop experimental setups to study their role in the absence of clinical antibiotics selection pressure.

Soft rot *Pectobacteriaceae* (SRP) are interesting model to analyze the role of β-lactam and β-lactamases in natural ecosystems. SRP are plant pathogens belonging to the genera *Pectobacterium* and *Dickeya* within the order *Enterobacterale*. SRP disrupt the plant cell wall and induce rotting symptoms on infected host plants (HugouvieuxlCottelPattat *et al*., 2014; Charkowski, 2018). SRP can infect important crops such as potatoes, causing a disease called blackleg in potato fields and a severe rotting of potato tubers called soft rot, which can lead to severe post-harvest losses. SRP act as a species complex and several species are often associated within symptoms (Degefu, 2021; de Werra *et al*., 2021; Ge *et al*., 2021; Motyka-Pomagruk *et al*., 2021; Barny *et al*., 2024). Recently, one species, *Pectobacterium versatile*, was described to harbor a β-lactamase, Bla_PEC-1_, that is the closet relative of Bla_TEM-1_, a class A2b β-lactamase plasmid-borne and widespread in clinical enterobacteria (Bush, 2018; Royer *et al*., 2022). The oldest identified *P. versatile* strain carrying the Bla_PEC-1_ β-lactamase was isolated from *Solanum tuberosum* in Canada in 1918, before the discovery of the first β-lactam antibiotic in the early 20th century ruling out the possibility that the acquisition of this gene within the chromosome is linked to the clinical use of β-lactam antibiotic (Royer *et al*., 2022). Phenotypic characterization *in vitro* revealed a low level of *P. versatile* Bla_PEC-1_ β-lactamase expression and it remains unclear whether Bla_PEC-1_ acts as a true β-lactamase during the infection process or whether it is merely a signaling molecules as previously proposed (Romero *et al*., 2011; Royer *et al*., 2022). Interestingly, within plant symptoms, *P. versatile* is often associated with other SRP species such as *Pectobacterium carotovorum* or *Pectobacterium brasiliense* and some strains of these latter species produce the structurally simplest member of the carbapenem class of β-lactam antibiotics, the (5R)-carbapen-2-em-3-carboxylic acid (Parker *et al*., 1982; McGowan *et al*., 1997, 2005). *In vitro*, this carbapenem is toxic to most *Pectobacterium* strains (Shyntum *et al*., 2019). However, carbapenem production is repressed in anaerobic conditions *in vitro* (McGowan *et al*., 2005; Shyntum *et al*., 2019). Furthermore, this carbapenem is highly unstable (Parker *et al*., 1982). Both the repression observed under anaerobic conditions and the instability of the carbapenem ring raise the question of carbapenem functional importance during potato infection.

The aim of the present work was to analyze the role of the *P. versatile* β-lactamase Bla_PEC-1_ in the context of plant infection when confronted with or not with *P. brasiliense* producing carbapenem. To this end, we constructed *bla*_PEC-1_-deleted strains in two different *P. versatile* strain backgrounds and compared them with their wild-type counterparts *in vitro* and *in planta,* with mono infection or mixed infections with different *P. brasiliense* strains.

## Experimental Procedures

### Bacterial strains, plasmids and growth conditions

The bacterial strains, plasmids and oligonucleotides used in this study are described in Tables S1 and S2 respectively. Briefly, the following strains have been used in the study: *P. brasiliense* type strain CFBP6617 (here after 6617; other names 1692, BAA-417, BPBB-212, Ecbr-212), *P. brasiliense* CFBP5381 (here after 5381), *P. versatile* type strain CFBP6051 (here after 6051; other names ICMP9168, NCPPB3387, de Boer 21) and its Δ*bla*_PEC-1_ mutant derivative CL1, *P. versatile* A73-S18-O15 (here after A73) and its Δ*bla*_PEC-1_ mutant derivative CL2, *E. coli* DH5-a-l-pir, *E. coli* MFD-pir. Strains were grown routinely at 30°C in LB (10 g/L tryptone, 5 g/L yeast extract, NaCl 5g/L), in TSB (1,7 g/L casein peptone, 0.3 g/L soya peptone, 0.3 g/L NaCl, 0.25 g/L dipotassium hydrogen phosphate, 0.25 g/L glucose) or in CVP (1.02 g/L CaCl_2_, 5 g tri-sodium citrate, 2.0 g/L NH_4_NO_3_, 4 g/L agar, 2.8 ml NaOH 5M, 18 g/L pectine Dipecta-AG366-Agdia-biofords,-USA) prepared as described by <u>Ben Moussa *et al*., 2023</u> for single layer CVP. When required, antibiotics were added at the following concentrations: ampicillin (Amp), 200 μg/L; Chloramphenicol (Cm), 50 μg/L.; Diaminopimelic acid (DAP) (57 μg/mL) was added for the growth of the *E. coli* MFD-pir strain. Media were solidified with 15 g/L agar.

### Construction of Δbla_pec-1_ mutants

To construct in-frame deletion mutants of the *bla*_pec-1_ gene, counter-selection method was used (Edwards *et al*., 1998). The cloning vector pRE112, that is an R6K-based suicide plasmid carrying the *sac*B gene and the *cat* gene (CmR) was used to clone into *Sac*I/*Kpn*I digested pRE112 two PCR fragments corresponding to the upstream and downstream 0.5-kbp DNA of the *bla*_pec-1_ gene using the T5 exonuclease-dependent assembly method (Xia *et al*., 2019).

Chemical ultracompetent DH5α λpir cells were prepared with the Mix & Go! *E. coli* Transformation Kit using standard procedures (Zymo Research). Transformants were selected onto LB plate supplemented with Cm. Colonies with the correct plasmid were selected by colony PCR with oligo pairs L762/L763 and DreamTaq DNA polymerase (Thermofisher, Waltham, MA, USA). Plasmids were extracted with the NucleoSpin Plasmid kit (Macherey-Nagel), checked by restriction digestion (NEB) and sequenced (Eurofins). Then, plasmids were transferred into competent *E. coli* strain MFDpir (Ferrières *et al*., 2010) prepared as described above for DH5α λpir. *E. coli* MFDpir produces the RP4 conjugation machinery, which allows the transfer of the suicide plasmid into *Pectobacterium* strains by conjugation. For conjugation, a few colonies of *P. versatile* A73 or *P. versatile* 6051 and were mixed in the same proportion with MFDpir colonies carrying the plasmid of interest in 500 µl LB and centrifuged for 2 min at 6000 rpm. The pellet was resuspended in 90 µl LB with 5 µl DAP at 57 mg/mL, and deposited onto a LB agar plate incubated at 28°C. After 24 h, the bacteria were resuspended in 1 ml LB, diluted in 10-fold series from 10-1 to 10-7 and spread onto LB agar supplemented with Cm at 4µg/l to select the first event of recombination. Transconjugants re-isolated on this medium were then spread onto LB agar supplemented with 5% sucrose and incubated at 19°C for 2-3 days to allow the second event of recombination. Sucrose-resistant colonies were then streak on LB-Cm plate to check plasmid loss and to LB-Amp plate to check for *bla*_pec-1_ gene loss. Total DNA of sucrose resistant, Cm and Amp sensitive colonies was used to amplified the mutagenized locus with the with the Prime start max DNA polymerase (Takara, kusatsu, Japan) and primers bla_pec-1_-sacI-up-fwd / bla_pec_-1-KpnI-down-rev and checked by sequencing (Eurofins). The resulting Δ*bla*_pec-1_ mutants constructed in strain *P. versatile* CFPB6051 and *P. versatile* A73 were named CL1 and CL2 respectively.

### In vitro growth inhibition assay

*Pectobacterium* strains were grown with shaking for overnight at 28°C in TSB medium. The next day, the optical density (OD_600nm_) of the culture was adjusted to 0.15. The temperature of the molten LB agar was lowered to around 40°C (just before the agar resolidified) and 25 mL of the LB agar were mixed with 1ml of the bacteria to be tested in lawn culture. A spot of 5 μl of strain 6617 was then added to the inoculated plates which were incubated at 30°C for 24–48 h before visualization of the inhibition zone.

### Potato tuber inoculations

Bacteria were plated on LB plates and incubated overnight at 28°C. A single colony was then used to inoculate 2 ml of LB medium, which was incubated overnight at 28°C with agitation at 120 rpm. This liquid culture (100 µl) was used to inoculate a 10 % TSB agar plate, which was incubated overnight at 28°C. The grown bacterial layer was then suspended in 50 mM phosphate buffer (pH 7) and adjusted to OD_600nm_=1. For co-inoculations, the strains involved were then mixed in equal volumes.

For each strain or strain mixture, 10 potato tubers were inoculated and each tuber was punctured with pipette tips containing 10 μl of the inoculum. The tip was left in the tuber. As a negative control, a pipette tip containing 10 µl of sterile phosphate buffer (50 mM) was inserted into a tuber. The inoculated tubers were placed on paper towels in plastic boxes, 50 mL of distilled water was poured into the bottom of the boxes and the boxes were carefully sealed to maintain relative humidity above 90%. After 5 days of incubation at 26°C, the tubers were cut and the entire macerating symptom was collected and weighed. To count the bacterial load in the symptom, part of the symptom (50 to 300 mg) was weighed and suspended in 1 ml of sterile phosphate buffer (50 mM) and diluted in 10-fold series and 100 µl of dilution 10^-5^ to 10^-7^ were spread on TSB plates and incubated 24h at 28°C. To analyzed the fitness of the Δ*bla*_pec-1_ mutant relative to the wild type strain, 100 individual colonies (per analyzed potato tuber) were streak on CVP, TSB-Amp and TSB agar plates. Colonies that formed pits on CVP, had the visual characteristics of *Pectobacterium* strains and grew on TSB-Amp agar plates were defined as *P. versatile* wild-type strains. Conversely, colonies that formed pits on CVP, had the visual characteristics of *Pectobacterium* strains and were unable to grow on TSB-Amp agar plates were defined as *P. versatile* Δ*bla*_pec-1_ mutant derivatives.

To recover the bacteria from rotten symptoms for DNA or RNA extraction, the rest of the symptom was suspended with 7 ml of 50 mM phosphate buffer and stirred for 4 hours at ambient temperature. The tubes were then left without shaking for 10 minutes to allow the excess starch to settle and 2 ml at the surface of the suspension was recovered and centrifuged for 10 minutes at 10,000 g. The formed pellet was kept at -80°C until DNA or RNA extraction.

### RNA extraction and RT-qPCR analysis

To investigate *carA* and *bla*_pec-1_ expression, RNA was isolated using PureLink® RNA Mini Kit from Thermo Scientific™, and 400 ng of RNA from potatoes symptoms or LB media were used to generate cDNA using the Thermo-scientific maxima First Strand cDNA Synthesis Kit for RT-qPCR with dsDNase. RT-qPCR experiments were performed with SsoAdvanced™ Universal SYBR® Green Supermix from BIO-RAD™. The housekeeping gene *gapA* was used to normalize the expression data for each gene of interest. The comparative quantitation method (ΔΔCt) was used to contrast the different treatments (Livak and Schmittgen, 2001). Ct values quantify the number of PCR cycles necessary to amplify a template to a chosen threshold concentration, ΔCt values quantify the difference in Ct values between a target gene and a the *gapA* gene for a given sample, and ΔΔCt values are used for the comparison between potato tuber symptoms and LB medium. Relative fold changes were calculated using 2^-ΔΔCt^.

### DNA extraction, DNA amplification and sequencing

DNA extraction was performed on 10 ml of the time zero inoculum, on 2 ml of 5 potato macerated tubers symptoms for each inoculated strain mixture. The bacteria were pelleted by centrifugation and DNA extraction was performed with the genomic DNA purification kit (Lucigen).

Amplification and sequencing were performed at MR DNA (www.mrdnalab.com, Shallowater, TX, USA). Either the *gap*A 376 partial gene sequence was amplified using PCR primers gapAF376 (GCCCGTCTCACAAAGA) and gapAR (TCRTACCARGAAACCAGTT) or the 16SrRNA gene V3-V4 region was amplified using PCR primers 341F (CCTACGGGNGGCWGCAG) and 805R (GGACTACHVGGGTWTCTAAT) with barcodes on the forward primer. PCR using the HotStarTaq Plus Master Mix Kit (Qiagen, USA) was performed under the following conditions: 94°C for 3 min, followed by 28 cycles of 94°C for 30 s, 57°C for 40 s (for *gapA* amplicon) or 53°C for 40s (for 16S amplicon) and 72°C for 1 min, after which a final elongation step at 72°C for 5 min was performed. Amplified PCR products were checked in 2 % agarose gel to determine the success of amplification and the relative intensity of bands. Multiple samples were pooled together in equal proportions based on their molecular weight and DNA concentrations. Pooled samples were purified using calibrated Ampure XP beads and used to prepare Illumina DNA library. Sequencing was performed on a MiSeq sequencer following the manufacturer’s guidelines. Sequence data were processed using MR DNA analysis pipeline (MR DNA, Shallowater, TX, USA). In summary, sequences were joined, depleted of barcodes and sequences <150bp or with ambiguous base calls were removed. Reads were filtered based on Q score and expected error probability and any read with a number of expected errors greater than 1.0 were discarded.

### Illumina sequencing analysis

After quality trimming, a total of 1,296,980 and 1,238,179 reads were obtained for 30 *gap*A and 20 16S analyzed potato tubers (50 samples *in toto*) assays respectively, with an average of 43,232 ± 8,490 and 61,908 ± 18049 reads per samples respectively.

In order to quantify the distribution of the *Pectobacterium* strains at time zero within the inoculum and 5 days after potato inoculation, SRP distribution analysis was performed at the strain level as previously described (Barny *et al*., 2024). Briefly, following *gap*A amplification, reads were aligned to the *gap*A sequences of the 6 strains analyzed (*P. brasiliense* 6617, *P. brasiliense* 5381, *P. versatile* A73 wild-type or Δ*bla*_PEC-1_ and *P. versatile* 6051 wild-type or Δ*bla*_PEC-1_), using the nucleotide-nucleotide Blastn tool (version 2.15.0+, e-value threshold 10^-5^). Only reads with 100% full-length identity were used (616,137 reads, 47.5% of the total reads).

The clustering, alignment and phylogenetic analysis of 16S gene fragments was performed using the QIIME2 (version 2023.5) (Bolyen *et al*., 2019) pipeline v.1.39.5, dedicated to the study of microbial communities. Demultiplexing of 16S rDNA gene sequences and quality control using DADA2 (Callahan *et al*., 2016). SILVA (version 138) (Bokulich *et al*., 2018) was used to perform taxonomic classification of each ASV. The feature table of ASVs was rarified to a sampling depth of 32,191 sequences per sample prior to downstream analyses.

## Results

### *In vitro*, the β-lactamase Bla_PEC-1_ allows *P. versatile* to resist to ampicillin or to the carbapenem produced by *P. brasiliense*

To analyze the role of Bla_PEC-1_, we constructed mutants deleted of the *bla*_PEC-1_ gene in two strains background of *P. versatile*, namely the type strain 6051 isolated from *Solanum tuberosum* and the strain A73 isolated from water. As expected, the two wild-type strains were resistant to ampicillin and the two Δ*bla*_PEC-1_ mutants became sensitive to this antibiotic, confirming that the ß-lactamase Bla_PEC-1_confers resistance to the ß-lactam ring of the penicillin class as previously observed by Royer *et al*. (2022) (Figure 1A). In contrast, the *P. brasiliense* 6617, which produces the carbapenem (5R)-carbapen-2-em-3-carboxylic acid, was sensitive to ampicillin. This sensitivity indicates that the *carF* and *carH* genes required for resistance to the carbapenem produced are inefficient against ampicillin (McGowan *et al*., 1997). We also tested another *P. brasiliense* strain, 5381, which was sensitive to both ampicillin and to the carbapenem produced by *P. brasiliense* 6617. When the *P. versatile* strains A73 or 6051 were confronted in competition assays with the carbapenem-producing strain *P. brasiliense* 6617, the two wild-type strains were resistant, while the two Δ*bla*_PEC-1_ mutants were sensitive, indicating that the β-lactamase Bla_PEC-1_ confers resistance to the simple carbapenem produced by *P. brasiliense* 6617 in addition to the penicillin class of ß-lactamase ring (Figure 1B).

**Figure 1:**
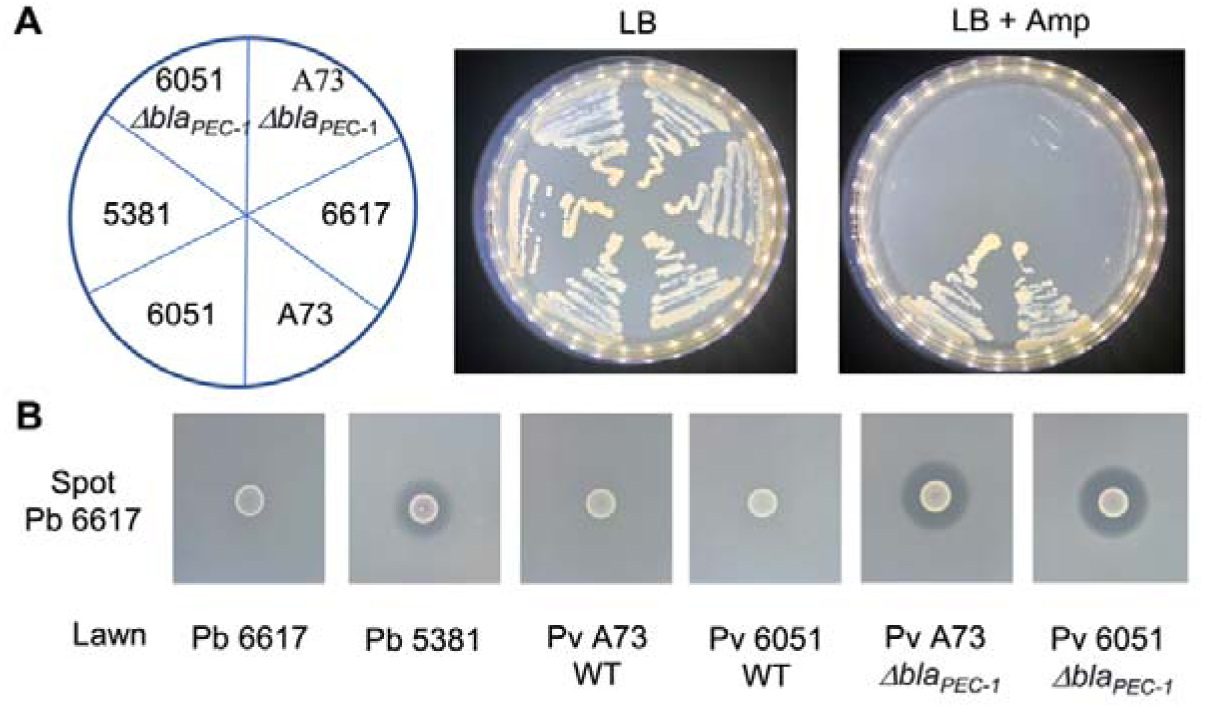
Spectrum of activity of Bla_PEC-1_ β-lactamase *in vitro*. A: Growth of the strains depicted in the left panel on LB and LB ampicillin (LB + Amp) media. *P. versatile* (Pv) strains: 6051 and A73; *P. brasiliense* (Pb) strains: 6617 and 5381. B: Growth inhibition assays between the carbapenem-producing strain Pb 6617 spotted and the lawn of strain Pb 6617, Pb 5381, Pb A73, Pv 6051, Pv A73 Δ*bla*_PEC-1_ mutant, Pv 6051 Δ*bla*_PEC-1_ mutant.

### In the absence of carbapenem, maceration capacity and fitness of *P. versatile* **Δ***bla*_PEC-1_ mutants in potato tubers are not altered compared to WT strains

We then investigated whether deletion of the *bla*_PEC-1_ gene altered the maceration capacity of the *P. versatile* strains in the absence of carbapenem. No difference in maceration capacity was observed between each wild-type strain and its Δ*bla*_PEC-1_ mutant derivative (Figure 2A). Strain A73 appeared to be slightly more aggressive than strain 6051, but this was only statistically significant when comparing the two wild-type strains. We then analyzed the fitness of each Δ*bla*_PEC-1_ mutant when co-inoculated with its wild-type counterpart (Figure 2B). The different susceptibilities to ampicillin of *P. versatile* wild type strains and their Δ*bla*_PEC-1_ mutant derivatives (see Figure 1A) were used to quantify the respective presence of each strain in potato tubers at the end of the co-infection experiment (Figure 2B). Within mix-inoculation, each wild-type strain and its Δ*bla*_PEC-1_ mutant derivative were present in the same proportion at the end of the experiment indicating that the β-lactamase Bla_PEC-1_ neither affects nor enhances the fitness of both strains in potato tubers.

**Figure 2:**
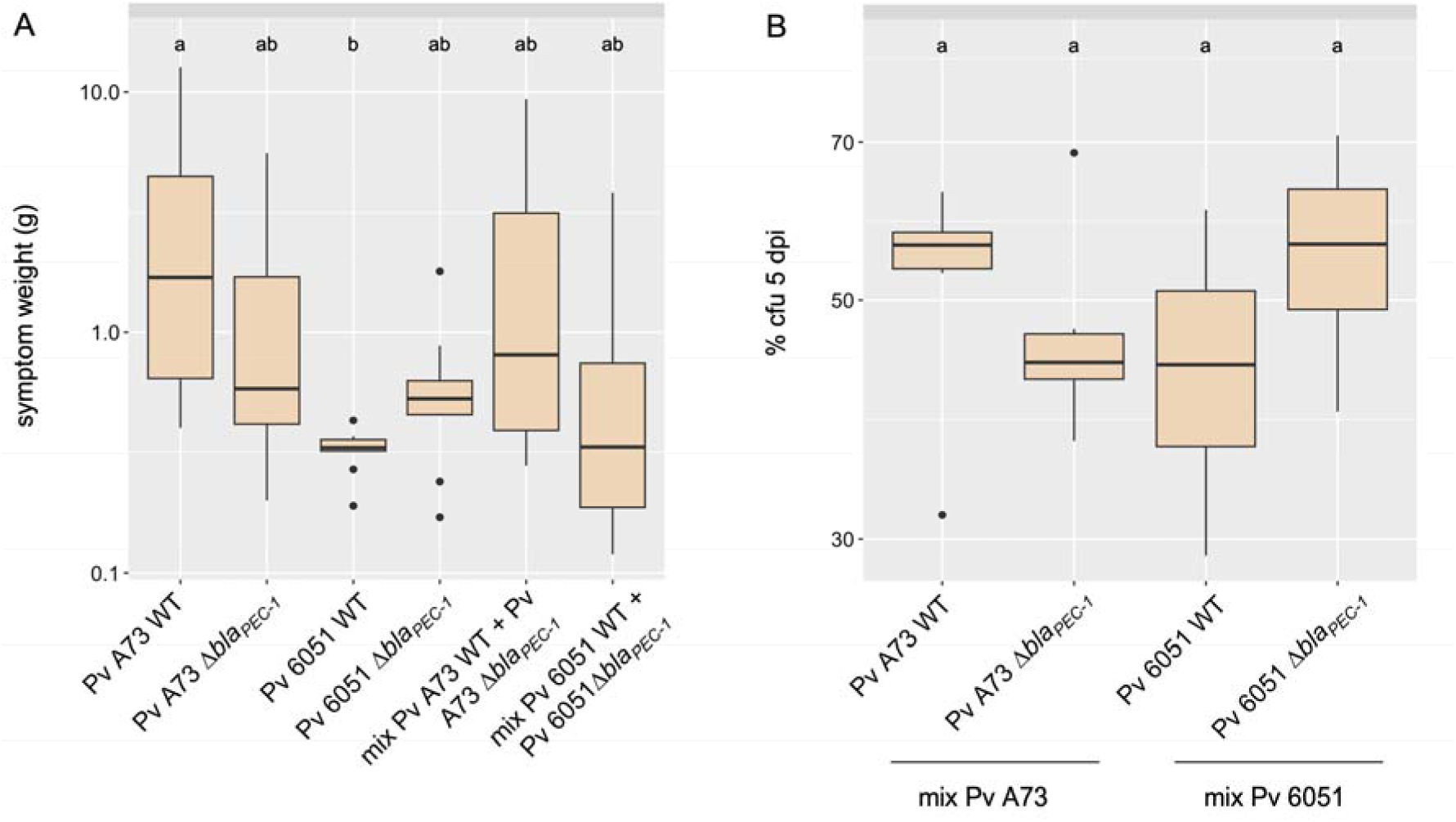
Maceration capacity and fitness of *P. versatile* Δ*bla*_PEC-1_ mutants. A: Weight of macerate symptom following inoculations of single species or mix infections. The species or mix that were inoculated on 10 potato tubers are indicated below each boxplot. B: The different susceptibilities of *P. versatile* 6051/A73 wild-type strains and 6051/A73 Δ*bla*_PEC-1_ mutant to ampicillin was used to assess the fate of each strain within mixed infections 5 days post infection (dpi). The boxplots represent the bacterial abundance of each strain at 5 dpi. The result of the statistical analysis is indicated by the letters (a, b,) at the top of the boxplots. The bacterial mixtures sharing the same letter(s) are not statistically different from each other (P>0.05 Kruskal Wallis followed by Dunn test with the Bonferroni correction).

### Coexistence of *P. versatile* wild-type strains with the carbapenem producing strain *P. brasiliense* 6617 in potato tuber

The resistance of the β-lactamase-producing wild-type strain A 73 and 6051 of *P. versatile in vitro* to the carbapenem produced by *P. brasiliense* 6617 prompted us to confront the strains *in planta*. To do this, we set up potato tuber infections with either single infection of each strain or co-infection between *P. brasiliense* and each of the wild-type *P. versatile* strains (10 tubers per condition). There was considerable variation in maceration capacity (Figure 3A). Statistical analysis showed that *P. brasiliense* strain 6617 was more aggressive than *P. versatile* 6051. Strain A73 showed an intermediate level of aggressiveness that was not statistically different from either 6051 or 6617. Co-inoculation of *P. brasiliense* 6617 with any *P. versatile* resulted in symptoms similar to single inoculation with 6617 or A73 suggesting that the mixed inoculation does not affect disease progression. The different susceptibilities of *P. brasiliense* 6617 and *P. versatile* 6051/A73 strains to ampicillin (see Figure 1A) were used to quantify the respective presence of each strain in potato tubers at the end of the co-infection experiment. After co-inoculation, *P. brasiliense* 6617 dominated the population at 5 dpi, consistent with its high level of virulence. Variability between inoculated tuber was important, however, both *P. versatile* strains were detected at 5 dpi in 8 out of 10 and 5 out of 10 inoculated potato tubers for A73 and 6051 respectively. Overall, the mean recovery of *P. versatile* A73 was 14,5% indicating its ability to co-exist with *P. brasiliense* 6617. In contrast, *P. versatile* 6051 was strongly outcompeted by strain *P. brasiliense* 6617 with a mean recovery of 1,2% (Figure 3B)

**Figure 3:**
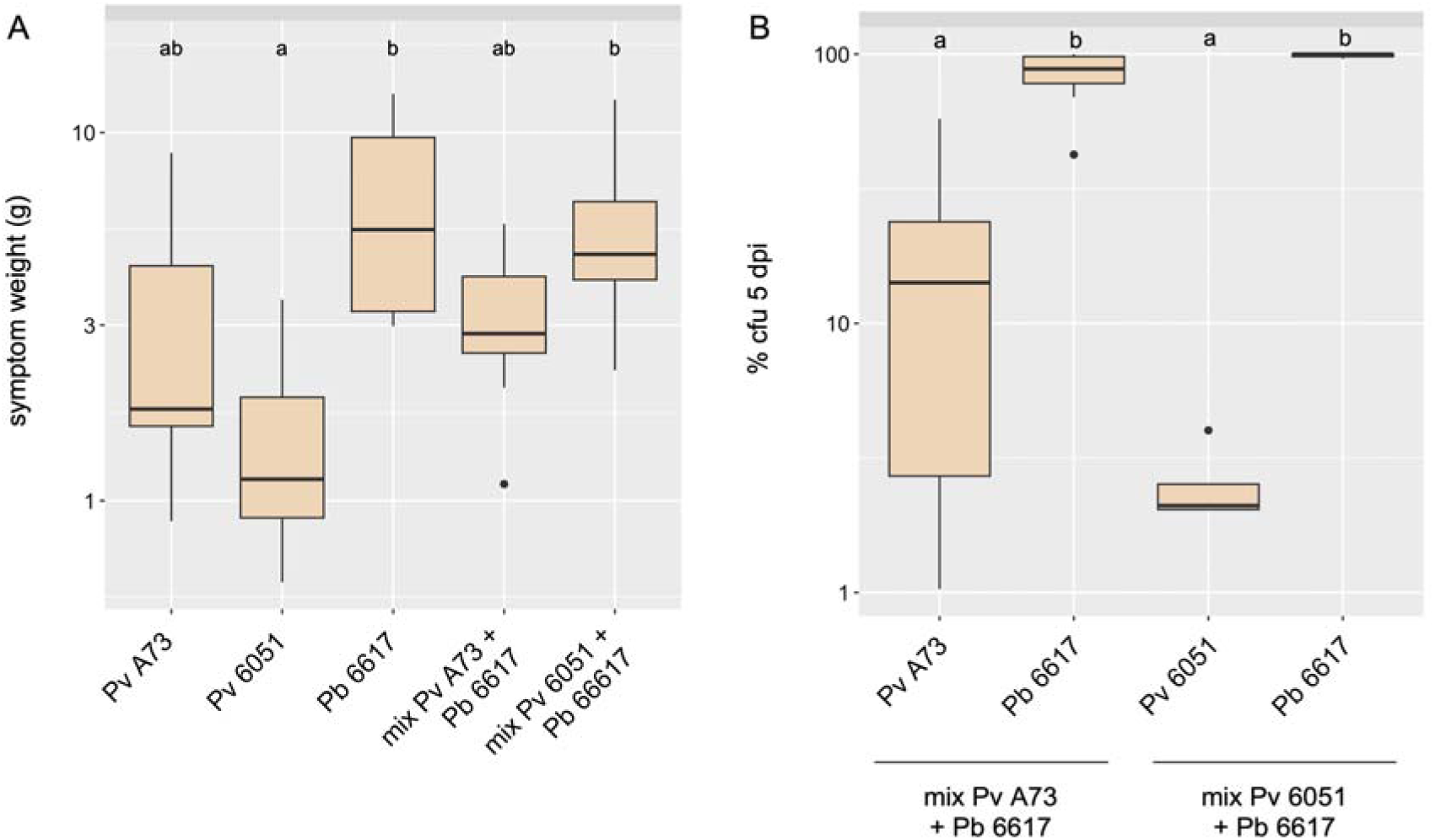
*In planta* competition between *P. versatile* strain expressing Bla_PEC-1_ and the *P. brasiliense* strain expressing carbapenem. A: Weight of macerate symptom following inoculations of single species or mix infections. The species or mix that were inoculated on 10 potato tubers are indicated below each boxplot. B: The different susceptibilities of *P. brasiliense* 6617 and *P. versatile* 6051/A73 wild-type strains to ampicillin was used to assess the fate of each strain within mixed infections 5 days post infection. The boxplots represent the bacterial abundance of each strain (% cfu 5 dpi). The result of the statistical analysis is indicated by the letters (a, b) at the top of the boxplots. The boxplots sharing the same letter(s) are not statistically different from each other (P>0.05 Kruskal Wallis followed by Dunn test with the Bonferroni correction).

### No association was found between commensal abundance in potato soft rot symptoms and the presence of the *P. versatile bla*_PEC-1_ gene

During infection of potato tubers, the important release of nutrients due to degradation of plant cell walls may allow the development of associated commensals <u>(Kõiv *et al*., 2015</u>). We tested whether the β-lactamase Bla_PEC-1_affects commensals invasion. Using 16S metabarcoding, we compared the abundance and diversity of commensals within symptoms at the end of the experiment between 20 tubers infected with either the wild-type strains or the Δ*bla*_PEC-1_ mutants (5 tubers per condition). The presence of commensals was often limited and was less than 1 % in 16 out of 20 tubers analyzed but a slightly higher proportion of associated commensals (from 2.1% to 14%) was observed in the 4 remaining potato tubers (Table 1). NMDS analysis indicated that commensal abundance was not statistically different between Δ*bla*_PEC-1_ mutants and their wild-type counterparts (Fig S1).

**Table 1:** Percentage of *Pectobacterium* and commensal at 5 dpi. Analysis was performed for each individual potato inoculated 5dpi (five replicates).

| Strain inoculated | potato | % *Pectobacterium* | % others |
| --- | --- | --- | --- |
| A73 WT | 1 | 99.6 | 0.4 |
|  | 2 | 99.6 | 0.4 |
|  | 3 | 99.1 | 0.9 |
|  | 4 | 99.6 | 0.4 |
|  | 5 | 99.7 | 0.3 |
| A73 $\Delta bla_{PEC-1}$ | 1 | 95.3 | 4.7 |
|  | 2 | 99.4 | 0.6 |
|  | 3 | 99.2 | 0.8 |
|  | 4 | 97.8 | 2.2 |
|  | 5 | 96.7 | 3.3 |
| 6051 WT | 1 | 99.7 | 0.3 |
|  | 2 | 99.7 | 0.3 |
|  | 3 | 99.6 | 0.4 |
|  | 4 | 99.7 | 0.3 |
|  | 5 | 86.1 | 13.9 |
| 6051 $\Delta bla_{PEC-1}$ | 1 | 99.7 | 0.3 |
|  | 2 | 99.6 | 0.4 |
|  | 3 | 99.4 | 0.6 |
|  | 4 | 99.7 | 0.3 |
|  | 5 | 99.6 | 0.4 |

We then analyzed the nature of the commensals recovered within the symptoms. The most common commensals to the 20 tubers analyzed were unclassified *Enterobacteriaceae* and bacteria belonging to the genus *Anaerocolumna* (Figure 4). Other commensals varied stochastically between potato tubers (Figure 4). In conclusion, the presence of commensals within symptoms was variable between tubers and did not correlate with the presence of the *bla*_PEC-1_ gene.

**Figure 4:**
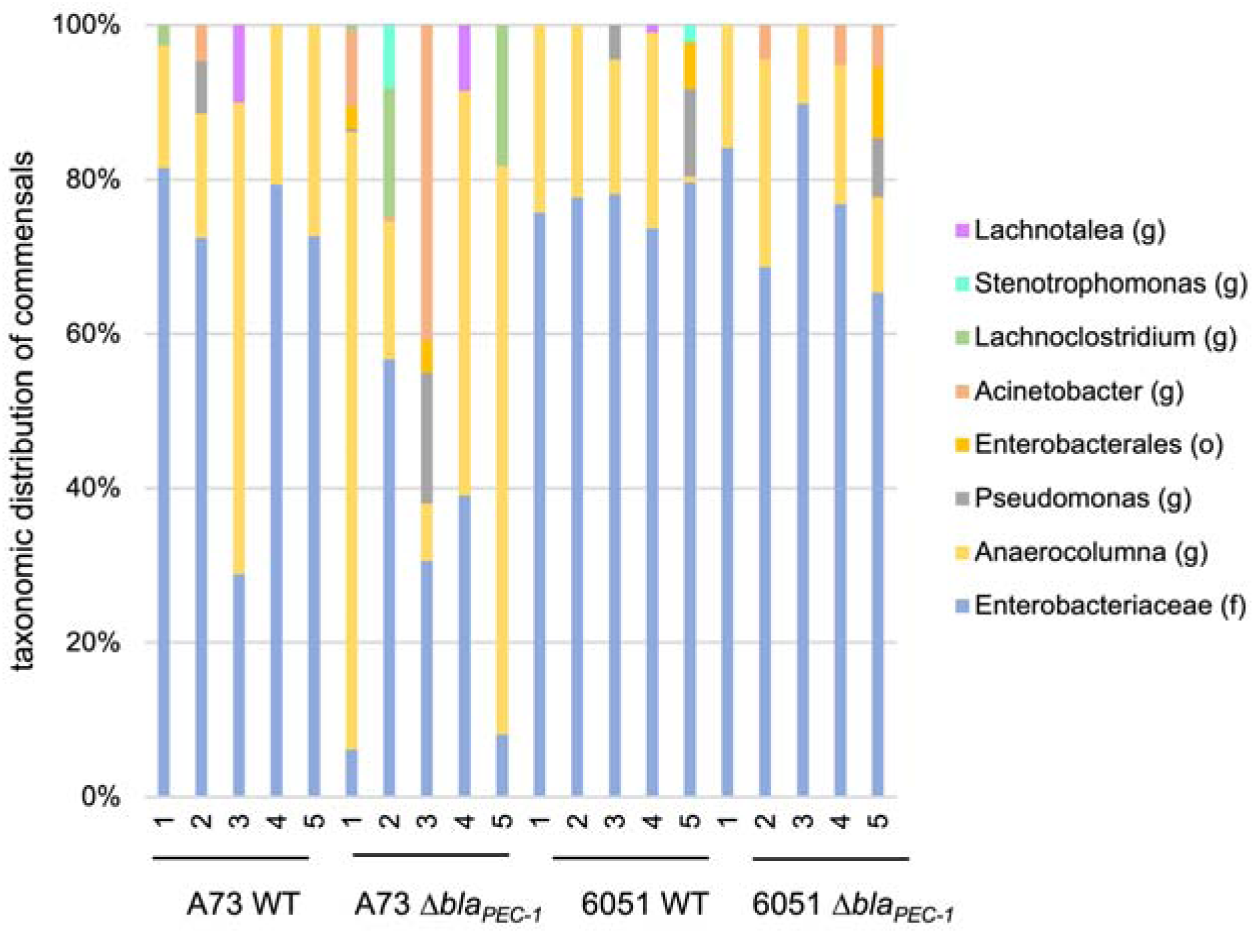
Commensal associated bacteria. Type of associated commensals in each analyzed potato tuber 5 days post infection following 16S barcode analysis. The inoculated strain and the potato number is indicated below each bar. The taxonomic assignment has been set at the genus level (g) or higher when this was not possible (f: family, o: order).

### Expression of carbapenem and **β**-lactamase encoding genes within potato tuber

To compared the level of expression of the genes involved in the synthesis of carbapenem or Bla_PEC-1_ β-lactamase during infection of potato tubers and LB synthetic media, we set up a mono-infection experiment with either the *P. brasiliense* strain 6617 or the *P. versatile* strains 6051 and A73. The expression levels of the *bla*_PEC-1_ and *carA* genes involved in the production of Bla_PEC-1_ β-lactamase and carbapenem respectively, were quantified by RT-PCR (McGowan *et al*., 1997; Royer *et al*., 2022). Large variations were observed between potato tubers. However, the *carA* gene of *P. brasiliense* 6617 was consistently strongly repressed within potato tubers as compared with LB media (Figure 5). Similarly, the *bla*_PEC-1_ gene was also repressed within potato tubers as compared with LB media but to a lesser extent than the one observed with the *carA* gene.

**Figure 5:**
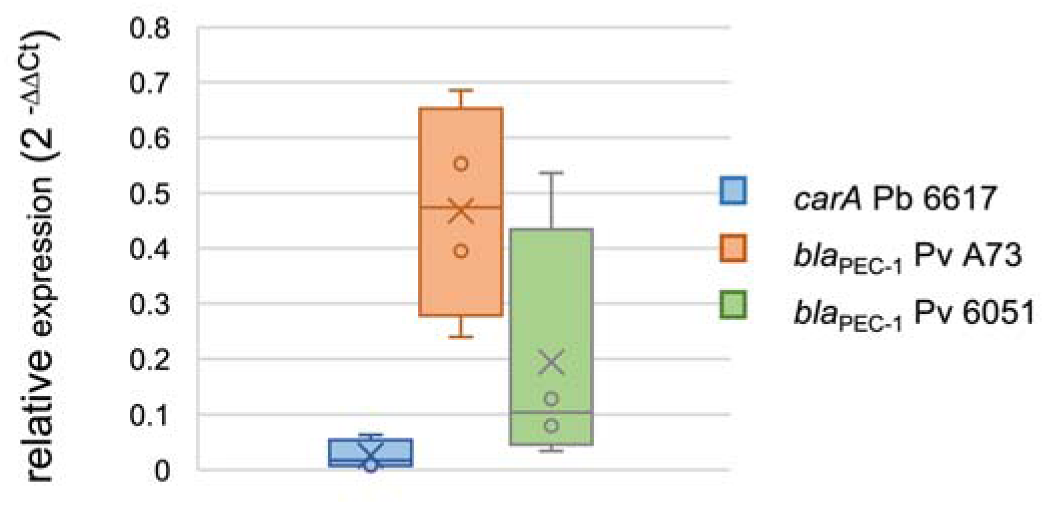
Relative expression of carbapenem and β-lactamase encoding genes within potato tuber. 2^-ΔΔCt^ values were used to express the fold change in *car*A (Pb 6617 strain) and *bla*_PEC-1_ (Pv A73 and 6051 strains) gene expression within potato tuber symptoms relative to expression in LB medium. A relative fold change of less than 1 indicates repression *in planta* compared to LB conditions *in vitro*. Expression data have been normalized using the housekeeping gene *gap*A as a reference gene.

### *P. versatile bla*_PEC-1_ gene is required during co-infection with carbapenem-producing *P. brasiliense* to protects another *P. brasiliense* strain sensitive to carbapenem

During infection, mixtures of SRP spp. can account for up to 20% of the observed symptoms (Degefu, 2021; de Werra *et al*., 2021; Ge *et al*., 2021; Motyka-Pomagruk *et al*., 2021). We therefore wondered whether the *P. versatile* β-lactamase Bla_PEC-1_ could protect other *Pectobacterium* strains from the carbapenem produced by the *P. brasiliense* strain 6617 (Shyntum *et al*., 2019). *In vitro*, the carbapenem-sensitive strain *P. brasiliense* 5381 was exposed to the carbapenem-producing strain 6617 in the vicinity of either the *P. versatile* β-lactamase-producing strain 6051 or its Δ*bla*_PEC-1_ mutants derivative. The presence of the *P. versatile* strain 6051, but not of the 6051 Δ*bla*_PEC-1_ mutant, protected the sensitive strain 381 from the toxic effect of *P. brasiliense* strain 6617 (Figure 6). This indicates that, *in vitro*, the β-lactamase Bla_PEC-1_ secreted by the WT strain of *P. versatile* is able to degrade the carbapenem produced and secreted by *P. brasiliense* 6617 efficiently enough to protect the sensitive strain 5381 when this latter strain is close enough to the β-lactamase PEC-1 produced by *P. versatile*.

**Figure 6:**
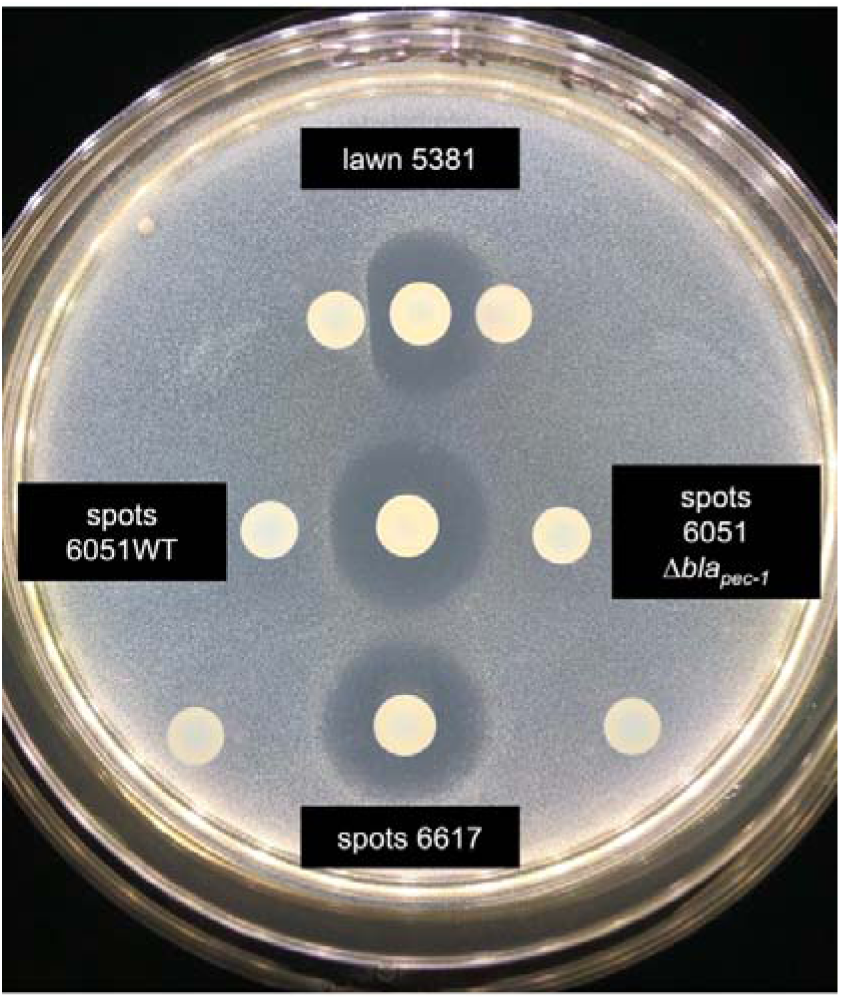
*In vitro*, the β-lactamase Bla_PEC-1_ confers protection against carbapenem. The *P. brasiliense* 5381 strain grown in the lawn is sensitive to the carbapenem produced by the *P. brasiliense* 6617 strain spotted in the middle of the Petri dish. The wild-type *P. versatile* strain 6051 spotted on the left, protects the sensitive strain only when the spot is in close proximity (compare upper and lower spots). None of the spots of the 6051 Dbla_PEC-1_ mutant of *P. versatile*, on the right, confers protection.

We then ask whether the protection conferred by *P. versatile* β-lactamase producing strains is also observed *in planta*. Potato tuber co-infection experiments were set up with 1) the *P. brasiliense* 6617 strain producing carbapenem, 2) the *P. brasiliense* 5381 strain sensitive to carbapenem and 3) either the wild-type *P. versatile* strains A73 or 6051 or their Δ*bla*_PEC-1_ mutant derivatives. As control, we also tested whether the carbapenem-sensitive strain 5381 is maintained when co-inoculated alone with the carbapenem producing strain 6617. The proportion of each strain at the beginning and end of the experiment was assessed by Illumina sequencing of the *gap*A discriminating gene marker (Barny *et al*., 2024). When the carbapenem-sensitive 5381 strain was co-inoculated alone with the carbapenem-producing strain 6617, it was outcompeted and the only strain recovered at the end of the experiment was the carbapenem-producing strain 6617 (Figure 7A). As well, in the mixture involving 6617 + 5381 and either the A73 Δ*bla*_PEC-1_ mutant or the 6051 Δ*bla*_PEC-1_ mutant, only the carbapenem-producing 6617 was recovered at the end of the experiment (Figure 7 B and C). In contrast, in the presence of the wild type *P. versatile* strains A73 or 6051, both the *P. versatile* strains and the sensitive strain 5381 were detect at various level the end of the experiment in 9 out of the 10 analyzed potato tubers (Figure 7 B and C). This indicated that the β-lactamase Bla_PEC-1_ protect both the *P. versatile* strain itself and the carbapenem sensitive strain 5381 from the toxic effect of the carbapenem produced by 6617 as observed *in vitro*. The protection conferred by strain 6051 was less than that conferred by strain A73, consistent with its weaker ability to compete with strain 6617. However, no correlation was observed between the proportion of *P. versatile* wild-type strains and the proportion of the carbapenem-sensitive strain 5381 recovered within the symptoms at 5dpi (Figure 7 B and C). This lack of correlation is likely explained by spatial effects within soft rot symptoms, as *in vitro* rescue of the carbapenem-sensitive strain 5381 strongly depends on spatial effects as observed *in vitro* (Figure 6).

**Figure 7:**
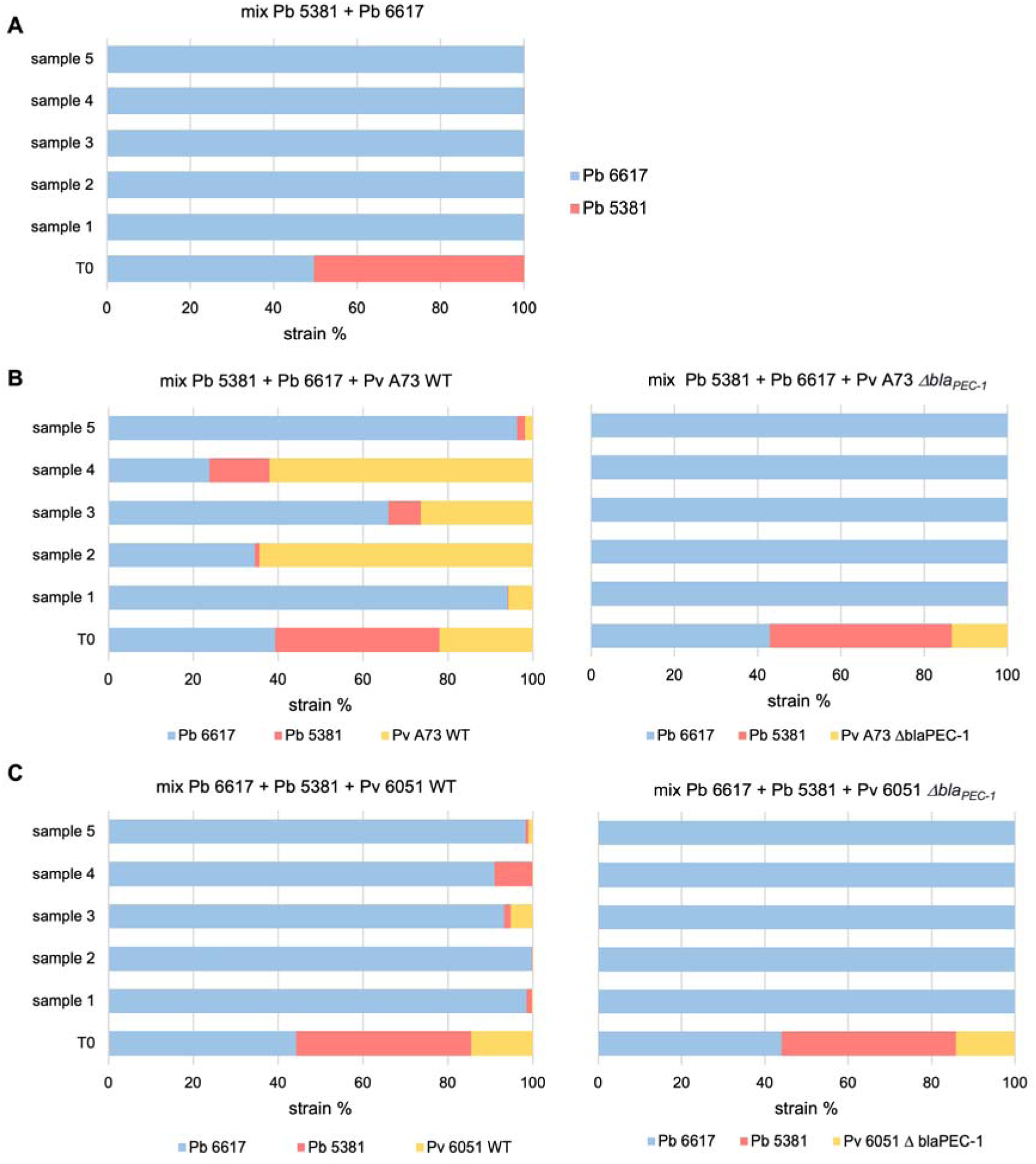
Fate of SRP consortium in mixed inoculation on potato tubers. The consortium inoculated (T0) is compared with the mix observed 5 days post inoculation in 5 independent potato tubers. The strains involved in each consortium are followed by analysis of the *gap*A barcode (Barny *et al*., 2024). Carbapenem producing strain are depicted in blue, carpapenem sensitive strain are depicted in red and β-lactamase producing strains are depicted in yellow. A: mix between the carbapenem producing *P. brasiliense* 6617 strain and the carbapenem sensitive *P. brasiliense* 5381 strain. B: mix between the carbapenem-producing *P. brasiliense* 6617 strain, the carbapenem-sensitive *P. brasiliense* 5381 strain and either the *P. versatile* A73 WT strain expressing the β-lactamase Bla_PEC-1_ or its Δ*bla*_PEC-1_ mutant derivative. C: mix between the carbapenem-producing *P. brasiliense* 6617 strain, the carbapenem-sensitive *P. brasiliense* 5381 strain and either the *P. versatile* 6051 WT strain expressing the β-lactamase Bla_PEC-1_ or its Δ*bla*_PEC-1_ mutant derivative.

## Discussion

The β-lactamase Bla_PEC-1_ is the closest relative of Bla_TEM-1_, a class A2b β-lactamase that is plasmid-borne and widespread in clinical enterobacterales. Bla_PEC-1_, like Bla_TEM-1_ cleave β-lactam rings of the penicillin family but is inefficient on clinically used carbapenem such as meropenem or imipenem (Royer *et al*., 2022). Here we have shown, using *in vitro* competition assays, that the β-lactamase Bla_PEC-1_ is nevertheless efficient in inactivating the simplest of the carbapenem molecule, produced by *P. brasiliense*. This suggests that the inability of Bla_PEC-1_ to inactivate imipenem and meropenem is probably due to the steric hindrance provided by the C-6 side chain of these clinically useful carbapenems. It remains to be determined whether the closest Bla_PEC-1_ β-lactamase, Bla_TEM-1_ that is clinically widespread around the world, is also able to cleave this simplest carbapenem molecule.

We then analyzed the role of the *P. versatile* β-lactamase Bla_PEC-1_ in the context of plant infection in the absence of antibiotic pressure. *In planta*, during mono-infection experiments, we could not detect a fitness cost associated with the presence of Bla_PEC-1_ as the wild-type and the *bla*_PEC-1_ -deleted strains multiplied similarly within symptoms. The fitness cost of a resistance determinant has been shown to partly depend on its genetic localization, such as a chromosomal mutation or a plasmid acquisition (Vanacker *et al*., 2023). The absence of fitness cost of associated with the *bla*_PEC-1_ gene could probably be partly due to the chromosomal location of this gene within the *P. versatile* genomes (Royer *et al*., 2022). It has also been shown that the fitness cost of antibiotic resistance genes is lower for β-lactamases compared to mutations in core genes which have a more pleiotropic effect that could be deleterious in the absence of antibiotic pressure (Vanacker *et al*., 2023). The absence of detectable fitness costs for chromosomally encoded β-lactamases in the absence of antibiotic pressure probably explains the wide distribution of these enzymes within natural communities. For example, in remote, undisturbed Alaskan soils with no antibiotic selection pressure, 5% of bacterial genomes carry diverse β-lactamase genes that are readily expressed in *Escherichia coli* (Allen *et al*., 2009). Similarly, permafrost bacteria represent a vast reservoir of antibiotic resistance genes, among which β-lactamase genes have been observed at high frequencies (Rigou *et al*., 2022).

Whether β-lactamases play a true role in antibiotic resistance in natural environments has been debated due to their low concentration (Davies *et al*., 2006; Aminov, 2009). *In vitro* phenotypic characterization of *P. versatile* revealed that Bla_PEC-1_ expression levels were low compared to that of Bla_TEM-1_ (Royer *et al*., 2022), suggesting that the role of Bla_PEC-1_ during infection could be merely a signaling role as previously proposed (Romero *et al*., 2011). Quantification of *bla*_PEC-1_ expression within the potato tuber showed that *bla*_PEC-1_ is repressed compared to its LB medium expression level. At the same time, the expression level of the *carA* gene, which is involved in the biosynthesis of the carbapenem by strain 6617, is also strongly repressed within the potato tuber. This is consistent with the dramatic repression of carbapenem expression observed *in vitro* in anaerobic medium (McGowan *et al*., 2005; Shyntum *et al*., 2019). Consistent with anaerobiosis, we found that commensals of the genus *Anaerocolumna*, a strictly anaerobic genus (Ueki et al., 2016), are systematically found associated with rotten symptoms in potato tubers. Surprisingly, despite the strong repression of *car*A in potato tubers, we found, that a carbapenem-sensitive strain was strongly outcompeted and could not be recovered when mixed inoculated with the carbapenem-producing strain. This indicates that carbapenem production is still efficient during infection, even though its expression is severely repressed compared to LB medium. The effectiveness of carbapenem in the potato tuber was also unexpected, as the carbapenem produced is also a highly unstable molecule. (Parker et al., 1982). This showed that lower expression level than the one observed *in vitro* and instability are not sufficient to rule out functionality and highlights the need for careful analysis to understand the effect of antibiotic production in natural environments.

When a *P. versatile* β-lactamase producing strain was added to the consortium, this third strain was able to rescue the carbapenem sensitive strain from the deleterious effect of the carbapenem producing strain. Such protection was space dependent and was observed *in vitro* only when the 3 strains were in close proximity. *In planta*, the protection was effective even when the *P. versatile* strain was a minority within the symptoms and the protection level, exemplified by the maintenance of the carbapenem sensitive strain, did not correlate with the proportion of each strain (protective, sensitive and antibiotic producing) within the symptoms. This likely highlights the formation of specific micro niches within the symptoms. This is in agreement with the fact that stochasticity in the initial phase of colony formation can be a crucial factor that determines the ensuing population dynamic (Von Bronk *et al*., 2017).

Within the SRP species complex, *P. versatile* occupies a special place as it is the most abundant SRP species, both on crops and in the environment, highlighting its ecological success (Portier *et al*., 2020; Ben Moussa *et al*., 2022). Among SRP species, *P. versatile* is also the only species that harbors the β-lactamase Bla_PEC-1_ at high frequency (Royer *et al*., 2022), suggesting that Bla_PEC-1_ may be involved in its ecological success. However, *P. versatile* is not considered to be a major pathogen of the SRP species complex, but rather a companion species during epidemics (van der Wolf *et al*., 2021). The work performed here indicates that Bla_PEC-1_ exerts a true β-lactamase function that is efficient at microscale during the infection process and highlights the role of *P. versatile* as a companion species that maintains strain diversity within the SRP species complex. The maintenance of such diversity is crucial to allow the regular emergence of new epidemic clones, such as those regularly observed on the potato host (Jonkheer *et al*., 2021). Beyond the SRP species complex, the high occurrence of various β-lactamases in natural pristine environments has recently been described (Lima-Bittencourt *et al*., 2007; Pedroso *et al*., 2017; Fonseca *et al*., 2018; Rigou *et al*., 2022). The work performed here unravels the role of β-lactamase in maintaining the diversity of β-lactam susceptible strains in the environment.

## Acknowledgments

Authors acknowledge Helena Dutey for her help in performing some plant experiments. The authors also acknowledge the funding from the ANR project ANR-19-CE35-0016-03. The funders had no role in study design, data collection and interpretation, or the decision to submit the work for publication.

## Data Accessibility

16S and *gap*A amplicon sequences are available at Zenodo (https://doi.org/10.5281/zenodo.14832342)

## Author contributions

M.A.B. and J.P. designed the research. E.G, M.A.B, and J.P. supervised research. C.L and P.Y.C performed the experiments. M.A.B and. J.P. analyzed the data. J.P. and P.Y. created the figures. M.A.B. wrote the first draft and all authors reviewed and edited the manuscript. M.A.B. was in charge of the funding acquisition.

## Supplementary Figure and tables

**Figure S1:**
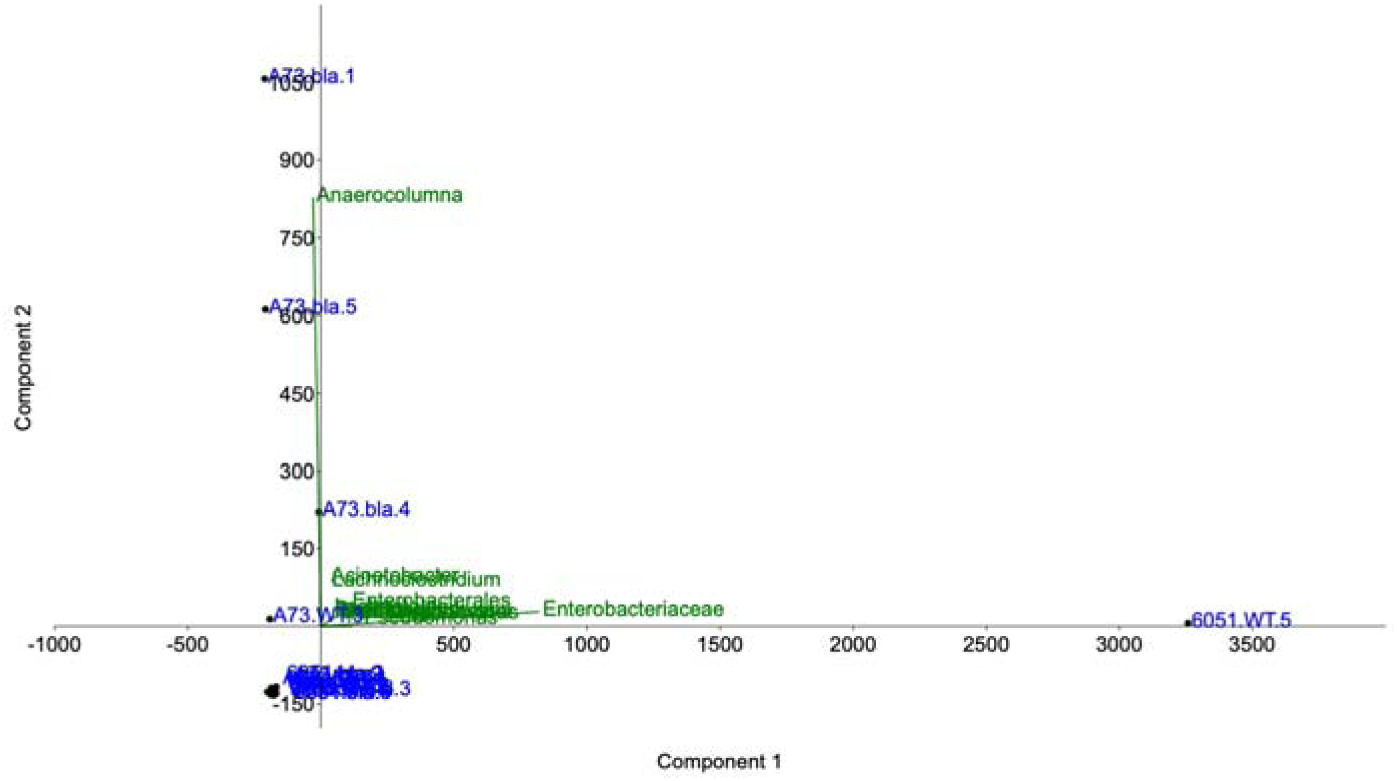
Principal components analysis of commensals. Principal component analysis was performed on the abundance matrix of commensal bacterial genera, 5 replicates per strain. Only 5 samples out of 20 (in blue) stand out, without forming a coherent group. For these samples, 2 variables (in green) are explanatory (genera *Anaerocolumna* and *Enterobacteriaceae*).

**Table S1:**
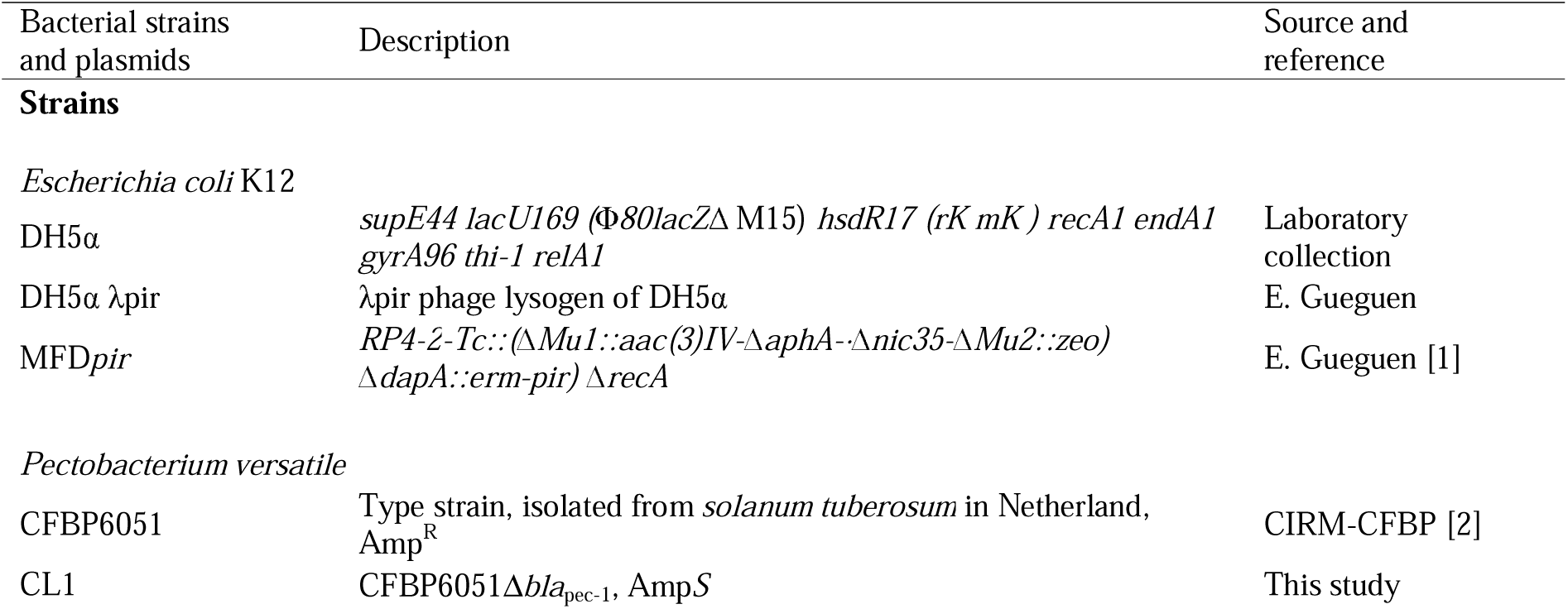
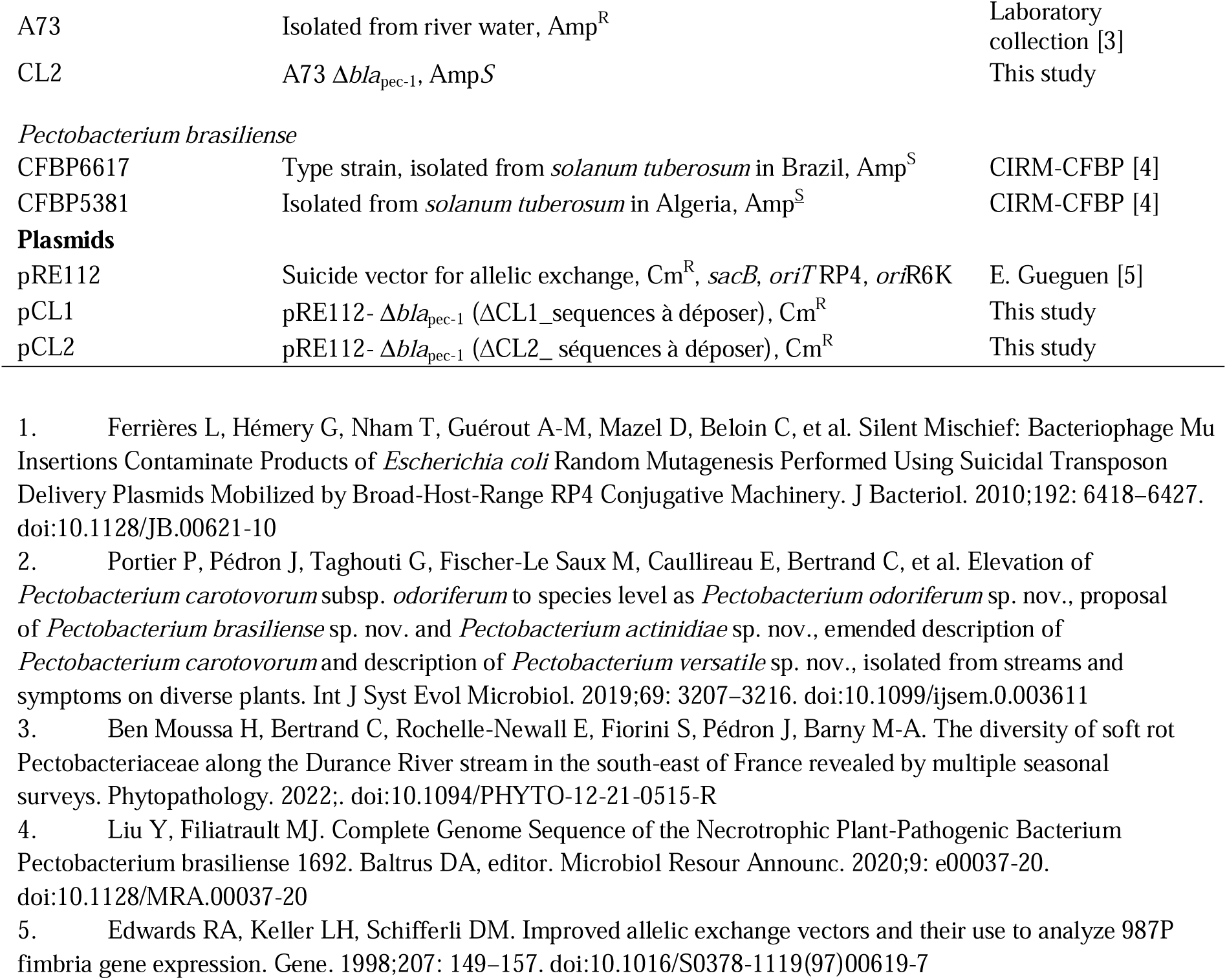
Bacterial strains and plasmids.

**Table S2:**
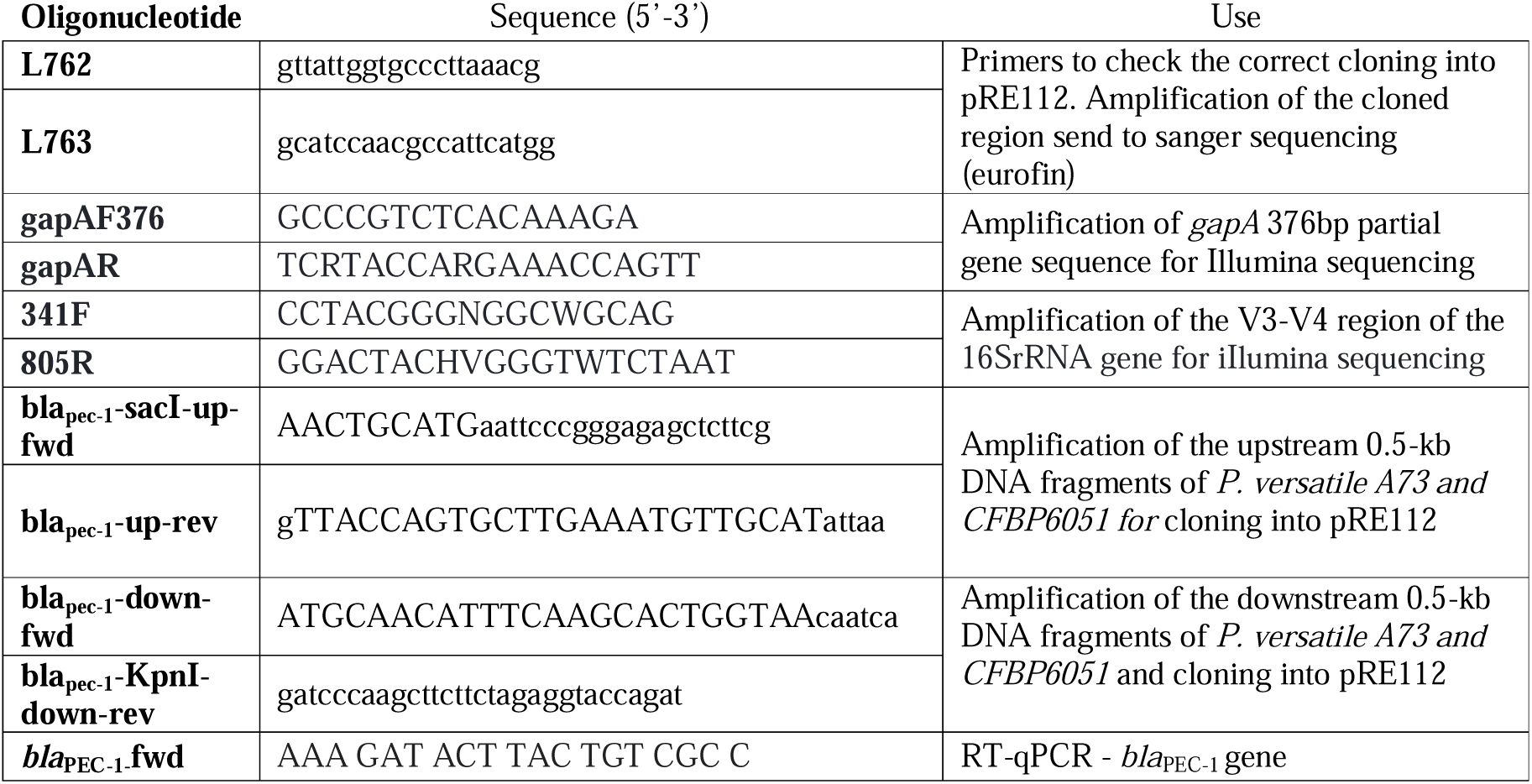
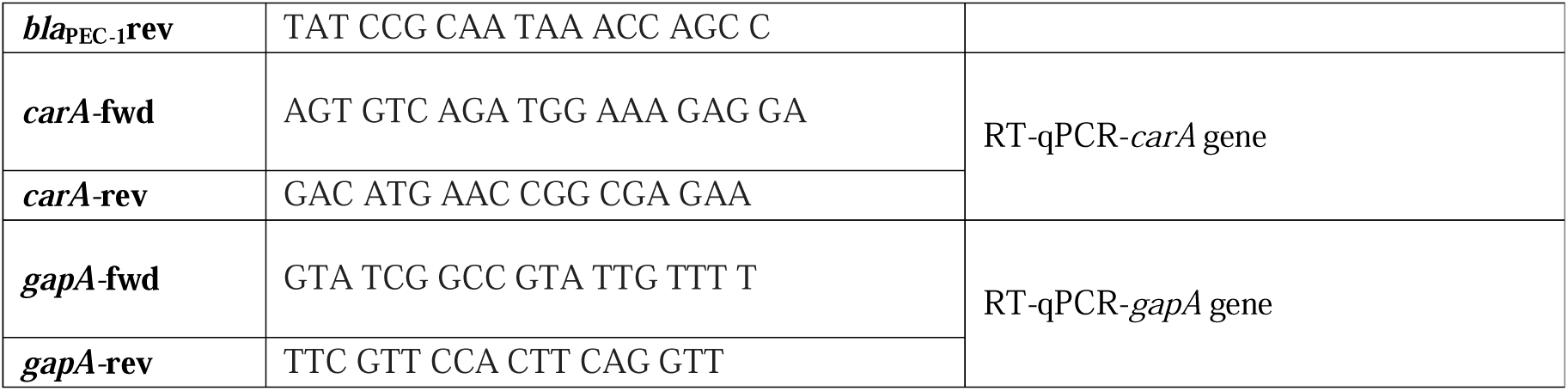
Oligonucleotides used in this study.

